# Experimental evolution of an RNA virus in *Caenorhabditis elegans*

**DOI:** 10.1101/2024.03.14.584948

**Authors:** Victoria G. Castiglioni, María J. Olmo-Uceda, Susana Martín, Marie-Anne Félix, Rubén González, Santiago F. Elena

**Affiliations:** Instituto de Biología Integrativa de Sistemas (CSIC-Universitat de València), Paterna, 46980 València, Spain; Institut de Biologie de l’École Normale Supérieure, CNRS, INSERM, 75005 Paris, France; Santa Fe Institute, Sant Fe, NM 87501, USA

## Abstract

The discovery of Orsay virus (OrV), the first virus infecting wild populations of *Caenorhabditis elegans*, has boosted studies of viral immunity pathways in this nematode. Considering the many advantages that *C. elegans* offers for fundamental research in host-pathogen interactions, this pathosystem has high potential to become a model system for experimental virus evolution studies. However, the evolutionary constraints operating in this pathosystem have barely been explored. Here we describe for the first time an evolution experiment of two different OrV strains in *C. elegans*. After 10 serial passages of evolution, we report slight changes in infectivity and non-synonymous mutations fixed in the evolved viral populations. In addition, we observed numerous minor variants emerging in the viral population. These minor variants were not randomly distributed along the genome but concentrated in polymorphic genomic regions. Overall, our work established the grounds for future experimental virus evolution studies using *Caenorhabditis* nematodes.

**HIGHLIGHTS:**

- *Caenorhabditis elegans*-Orsay virus is a convenient pathosystem to study virus evolution.
- The approach used to test the viral strains may interfere with the infection phenotypes observed.
- There may be specific genomic hotspots regions of nucleotide diversity important for the evolution of Orsay virus.
- The substitution rate observed for Orsay virus was low, suggesting that the two strains studied might be already well adapted to laboratory conditions.

## INTRODUCTION

The experimental evolution of microbes in their hosts enables the study of evolution in real time using controlled settings. Evolution experiments deepen our understanding of evolutionary principles and the evolution of specific traits, and are useful for the validation of evolutionary theories and the discovery of novel phenomena (Ebert, 1998; Elena and Lenski, 2003). Additionally, apart from providing insights into parasite evolution, experimental evolution has practical applications by enabling the directed evolution of microbes for industrial and medical purposes (McDonald, 2019; Lewis and Morran, 2022).

Within the microbial world, viruses are recognized as complex adaptive systems with high evolvability (Solé and Elena, 2019). Their high mutation rates, which vary based on the nucleic acid composition and size of their genomes (Sanjuán, 2010), lead to the formation of highly polymorphic populations with a particular genetic structure known as viral quasispecies. Viral quasispecies consist of a swarm of low frequency variants that continually emerge around a dominant, master sequence (Domingo and Perales, 2019). Despite the potential for deleterious mutations in their compact genomes, the large population sizes and the quasispecies structure provide a safeguard and ensure the persistence and adaptation of the most fit variants at the population level (Elena, 2012). It is precisely this interplay between high mutation rates, compact genomes, and large, diverse populations what renders viruses particularly well-suited for evolutionary studies (Elena and Sanjuán, 2007).

Viruses have been experimentally evolved in widely diverse hosts, such as bacteria (Bull et al., 1997; Wichman et al., 1999; Meyer et al., 2012), cell cultures (Elena et al., 1996; Keleta et al., 2008; Poole et al., 2014; Garijo et al., 2014; Grass et al., 2022), wild birds (Grubaugh et al., 2015), mice (Kubinak et al., 2013, Cornwall et al., 2018), and plants (González et al., 2021; Navarro et al., 2022). The use of model organisms with convenient experimental characteristics has expanded the repertoire of environmental and genetic conditions that can be manipulated during experimental evolutions. Examples are the plant *Arabidopsis thaliana* (Pagán et al., 2010; Elena, 2017) and the insect *Drosophila melanogaster* (Martínez et al., 2019; Mongelli et al., 2022; Lezcano et al., 2023). However, despite its many advantages for dissecting the genetic and molecular basis of host-pathogen interactions, *Caenorhabditis elegans* MAUPAS has not been used for experimental virus evolution yet.

*C. elegans* is an easy to maintain nematode with a powerful genetic toolbox available (Brenner, 1974; Nance and Frøkjær-Jensen, 2019). *C. elegans* lacks an adaptive immune system but defends itself against pathogens through innate and nonclassical adaptive immune responses (Martineau et al., 2021; Tran and Luallen, 2023; González and Félix, 2024a). Orsay virus (OrV) was the first natural virus isolated in *C. elegans* (Félix et al., 2011). In addition to OrV, multiple viruses that infect nematode species closely related to *C. elegans* have been found (Frézal et al., 2019; Richaud et al., 2019), but OrV is still the only natural virus that infects *C. elegan*s (Félix and Wang, 2019). OrV is a positive sense single-stranded RNA virus similar to nodaviruses. OrV has a bipartite genome: RNA1 encodes for the RNA-dependent RNA polymerase (RdRp), and RNA2 encodes for the capsid (CP) and δ proteins as well as a CP-δ fusion protein, the latter produced through a ribosomal frameshift (Jiang et al., 2014; Félix and Wang, 2019). OrV is orofecally transmitted between individuals: new hosts ingest virions while feeding, the virus replicates in a few intestinal cells where new infectious virions are assembled, exit without disruption of the host cell and are released outside the host (Yuan et al., 2018; Félix and Wang, 2019). Infected animals do not have a reduced lifespan but have a reduced brood size and their progeny production is delayed (Félix et al., 2011; Ashe et al., 2013). OrV was originally isolated from the *C. elegans* isolate JU1580 sampled in Orsay, France (Félix et al., 2011) and its infecting viral strain, JUv1580, has been widely used in subsequent studies. Later on, another OrV strain, JUv2572, was isolated from *C. elegans* JU2572 at Ivry, France. Comparing the infectivity of different OrV strains can be cumbersome due to (*i*) the usually low infectivity that introduces noise in dose-response curves, and (*ii*) quantifying viral load by relative RT-qPCR faces limitations in accurately reflecting the actual number of infectious viral particles. To overcome these difficulties, Frézal et al. (2019) stablished a commonly used protocol based in single infections and testing the number of infected animals in the population over two to three host generations, ensuring that the observed outcomes are due to host-pathogen interactions rather than differences in the initial inocula. Using this approach, the JUv2572 viral strain has been described to be more infectious and to infect more anterior cells than JUv1580 (Frézal et al., 2019).

In this study, our aim was to evolve OrV JUv1580 and JUv2572 strains in *C. elegans*. We described phenotypic changes in experimentally-evolved OrV lineages as well as hotspots for nucleotide diversity along the genome. Our work paves the path for using the *C. elegans*-OrV pathosystem in future experimental evolution research.

## MATERIAL AND METHODS

### *C. elegans* strain maintenance

Nematodes were maintained at 20 °C on Nematode Growth Medium (NGM) plates seeded with *Escherichia coli* OP50 under standard conditions (Brenner, 1974; Stiernagle, 2006). ERT54 (*jyIs8 [pals-5p::GFP + myo-2p::mCherry]X*) (Bakowski et al., 2014), derived from N2, was used in all experiments except for ancestral viral stock preparation, for which JU2624 (*[myo-2p::mcherry::unc-54; lys-3p::eGFP::tbb-2] IV*), derived from JU1580 background, was used instead.

### OrV stock preparation and quantification

For ancestral JUv1580_vlc or JUv2572_vlc stock preparation (we named the strains “vlc” as they were characterized in Valencia, Spain, and to distinguish them from the original field isolates), JU2624 animals were inoculated with either JUv1580 or JUv2572 (Félix et al., 2011; Frézal et al., 2019). For evolved virus stock preparation, ERT54 animals were inoculated with the corresponding evolved stocks. In either case, animals were allowed to grow for 5 days and then resuspended in M9 (0.22 M KH_2_PO_4_, 0.42 M Na_2_HPO_4_, 0.85 M NaCl, 0.001 M MgSO_4_), let stand for 15 min at room temperature, vortexed, and centrifuged for 2 min at 400 g. The supernatant was centrifuged twice at 21,000 g for 5 min and then passed through a 0.2 μm filter. RNA of the resulting viral stock was extracted using the Viral RNA Isolation kit (NZYTech). The concentration of viral RNA was then determined by RT-qPCR using a standard curve, and normalized across different stocks (details below). Primers used for RT-qPCRs can be found in Table S1.

For the standard curve, cDNA of JUv1580_vlc was obtained using AccuScript High-Fidelity Reverse Transcriptase (Agilent) and reverse primers at the 3’ end of the genome. Approximately 1000 bp of the 3’ end of RNA1 and RNA2 were amplified using forward primers containing 20 bp coding the T7 promoter and DreamTaq DNA Polymerase (Thermo Fisher). The PCR products were gel purified using MSB Spin PCRapace (Invitek Molecular) and an *in vitro* transcription was performed using T7 Polymerase (Merck). The remaining DNA was then degraded using DNAse I (Life Technologies). RNA concentration was determined by NanoDrop (Thermo Fisher) and the number of molecules per µL was determined using the online tool EndMemo RNA Copy Number Calculator (https://www.endmemo.com/bio/dnacopynum.php). Primers used for the standard curve can be found in Table S1.

### Virus inoculation

In order to obtain synchronized animal populations, plates with embryos were carefully washed with M9 buffer to remove larvae and adults but leaving the embryos behind. Plates were washed again using M9 buffer after 1 h to collect larvae hatched within that time span. Synchronized populations were inoculated by pipetting the viral stock on top of the bacterial lawn containing the animals. The normalized inoculum had 3.6×10^7^ copies of OrV RNA2. Three-hundred animals per plate were grown for RT-qPCRs and fluorescence *in situ* hybridization (FISH).

### RNA extractions and RT-qPCRs

RNA extractions were performed in two different ways. For RT-qPCRs and sequencing of the 3’ ends of JUv2572_vlc, animals were collected with PBS-0.05% Tween, centrifuged for 2 min at 1350 rpm and the supernatant discarded. Two more wash steps were performed before freezing the samples in liquid N_2_. Five-hundred µL of Trizol (Invitrogen) were added and the pellet was disrupted by following five cycles of freeze-thawing and five cycles of 30 s of vortex followed by 30 s of rest. One-hundred µL of chloroform were then added and the tubes were shaken for 15 s and let rest for 2 min. Samples were centrifuged for 15 min at 11,000 g at 4 °C and the top layer containing the RNA was then mixed with the same volume of 100% ethanol. Samples were then loaded into RNA Clean & Concentrator Kit columns (Zymo Research) and the rest of the protocol was followed according to manufacturer instructions.

RT-qPCRs were performed using Power SYBR Green PCR Master Mix (Applied Biosystems) on an ABI StepOne Plus Real-time PCR System (Applied Biosystems). Ten ng of total RNA were loaded and a standard curve was used to quantify OrV. Primers used for RT-qPCRs can be found in Table S1.

Five biological replicates per viral strain were evaluated, with 300 animals per replicate. A generalized linear model (GLM) with a gamma distribution and a log-link function was employed to evaluate the impact of the viral strains and viral segments on the RNA accumulation. Viral strain and segment were treated as fixed orthogonal factors, while biological replicate was a random factor nested within the interaction of the two fixed factors.

For sequencing the genome and 5’ ends of JUv2572_vlc RNA was extracted using the Viral RNA Isolation Kit (NZYTech) following manufacturer instructions.

### FISH

The protocol was adapted from previous studies (Parker et al., 2021; Tsanov et al., 2016). Virus-inoculated and mock-treated nematodes were washed with PBS-0.05% Tween and centrifuged for 2 min at 1350 rpm four times discarding the supernatants. Eight hundred µL of Bouins Fixation mix (400 µL Bouins Fix, 400 µL methanol, 10 µL β-mercaptoethanol) were added and the sample incubated at room temperature for 30 min in a rotatory shaker. Samples were then frozen in liquid N_2_ and kept at −80 °C overnight. The following day, samples were gently shaken at 4 °C for 30 min and washed four times with borate Triton solution (20 mM H_3_BO_3_, 10 mM NaOH, 0.5% Triton) and five times with borate Triton β-mercaptoethanol solution (20 mM H_3_BO_3_, 10 mM NaOH, 0.5% Triton, 2% β-mercaptoethanol), leaving the borate Triton β-mercaptoethanol incubate for 1 h between each wash in the last three washes. The sample was then incubated for 5 min with Wash Buffer A (1 mL Stellaris RNA FISH Wash Buffer A, 1 mL deionized formamide and 3 mL H_2_O).

Forty probes against OrV were designed using Oligostan (Tsanov et al., 2016): 22 against RNA1 and 18 against RNA2. The sequences of the probes can be found in Table S1. The probes were annealed as follows: 2 µL 0.833 µM probe set, 1 µL 50 µM FLAP-label, 1 µL NEB3 and 6 µL DEPC-H_2_O were incubated at 85 °C for 3 min, 65 °C for 3 min and 25 °C for 5 min. The FLAP labels contained CAL Fluor 610 or Quasar 670 modifications at both the 5’ and 3’ ends of the following sequence: ^5’^AATGCATGTCGACGAGGTCCGAGTGT_3’_ (Biosearch Technologies). One µL of annealed probe solution targeting RNA1 and 1 µL of annealed probe solution targeting RNA2 was then mixed with 98 µL of Hybridization Buffer (100 µL Stellaris FISH hybridization buffer, 25 µL deionized formamide). One hundred µL of Hybridization Buffer containing multiple FISH probes were then added to the samples and the samples incubated overnight at 37 °C in a rotary shaker. The following day, samples were washed with Wash Buffer A2 (1 mL, Stellaris RNA FISH Wash Buffer A, 4 mL H_2_O), incubated for 30 min in Wash Buffer A2 at 37 °C, and then incubated for 30 min in Wash Buffer A2 containing 25 ng of DAPI at 37 °C. Samples were then incubated 5 min in Stellaris RNA FISH Wash Buffer B, centrifuged and resuspended in a small volume of Stellaris RNA FISH Wash Buffer B. DAPI (0.1 ng) were added to each sample and mounted using N-propyl gallate mounting medium. Samples were imaged using Leica DMi8 microscope with Leica DFC9000 GTC sCMOS camera and objective HC PL APO 40×/0.95 CORR PH2. Images were analyzed and processed using ImageJ (FIJI) (Schindelin et al., 2012). FISH signal was used to determine the number of infected cells in each animal and the localization of the signal (*i.e.*, within intestinal cells, in the intestinal lumen or both).

Four populations of 50 animals each were evaluated per viral strain. The counts of infected animals were fitted to a GLM with a binomial distribution and a logit-link function to evaluate the impact of the viral strains on the proportion of infected animals. The model included viral strain as main fixed factor and replicate nematode population as random nested factor. For each of the infected animals, the number of FISH positive cells was also recorded. These counts were fitted to GLM with a Poisson distribution and a log-link function with the same structure as in the previous analysis. Finally, the counts of infected animals were fitted to a second GLM with a binomial distribution and a logit-link function but now adding as additional fixed main factor the localization of the FISH signal. In this model, viral strain and localization were treated as orthogonal and nematode population nested within their interaction.

### Fluorescent reporters for viral infection

The *pals5p::GFP* signal on the animals was visualized under the fluorescent dissecting microscope and the detection of the signal was used to determine the activation of response to infection. From the total number of animals analyzed, the percentage of infected animals was calculated. The infection status was determined either at 24 hours post-inoculation (hpi) when using normalized viral stocks or after performing three consecutive passages in which nematodes were allowed to develop for 5 d before transferring 20 or 50 animals at developmental stage L4 to a new plate. After the three passages, L4s were placed in a new plate and the presence or absence of the *pals5p::GFP* signal was quantified.

Five replicate populations were evaluated per condition, with 50 animals assayed per replicate. A GLM with a binomial distribution and a logit-link function was fitted to the counts of GFP positive animals. The model incorporated viral strain and the passage protocol described above as orthogonal main factors. Replicate population was treated as a random factor nested within the interaction of the two main factors.

### Experimental evolution of OrV

Ten ERT54 larvae at developmental stage L1 were inoculated with JUv1580_vlc or JUv2572_vlc and the population expanded for 6 d. If activation of the *pals-5p::GFP* signal was visible under the fluorescent binocular, nematodes were resuspended in M9, incubated for 5 min at room temperature, vortexed and centrifuged for 2 min at 400 g. The supernatant was centrifuged twice at 21,000 g for 5 min and then filtered through 0.2 μm. The resulting viral preparation was used to inoculate fresh L1 larvae and the process was repeated 10 times in triplicate. The evolution of JUv2572_vlc was replicated in two fully independent blocks, hereafter, referred to as V and R. After the 10^th^ passage, viral stocks were prepared and RNAs extracted for stock normalization using Viral RNA Isolation kit (NZYTech) and for RNA-seq using Trizol (Invitrogen), as detailed above.

The number of GFP positive animals was fitted to a GLM with a binomial distribution and a logit-link function, where the main orthogonal factors were the original strain from which the virus evolved and the ancestral/evolved status of the replicate viral population. Replicate populations were nested within the interaction of the two main factors. To test the consistency between the two blocks of evolution experiments, another GLM was fitted to the counts of positive animals infected with JUv2572-derived R and V lineages, using block as main factor and the number of replicated populations as nested factor, both taken as random.

### Viral genome sequencing

To obtain the genome sequence of the JUv1580_vlc and JUv2572_vlc, cDNA of OrV RNA1 was obtained using SuperScript III Reverse Transcriptase (Invitrogen) and primer oVG6_OrV_RNA1_3’_R cDNA of OrV RNA2 was obtained using AccuScript Reverse Transcriptase (Agilent) and primer oVG61_OrV_RNA2_R cDNA complementary to 3’ ends were synthesized from negative viral strands using forward primers. cDNAs were amplified using Phusion High-Fidelity DNA polymerase (Thermo Fisher) and primers oVG6_OrV_RNA1_3’_R and oVG75_OrV_RNA1_F, and oVG61_OrV_RNA2_R and oVG60_OrV_RNA2_F, for RNA1 and RNA 2, respectively. PCR products were purified using Zymoclean™ Gel DNA Recovery Kit (Zymo Research) and sequenced by Sanger. Viral 5’ terminal positions were determined using the 5’RACE System for Rapid Amplification of cDNA Ends (Invitrogen). To obtain 3’ ends, cDNAs were synthesized from negative viral strands using forward primers, and the rest of procedure was performed according manufacture’s instructions. Primers used can be found in Table S1.

### High-throughput sequencing of evolved strains

Library preparation and Illumina sequencing was done by Novogene Europe using a NovaSeq PE150 platform and a Lnc-stranded mRNA-Seq library method, ribosomal RNA depletion and directional library preparation, with 150 paired-end reads and 6 GB raw data per sample. The quality of the libraries was checked using a Qubit 4 Fluorometer (Thermo Fisher) and Bioanalyzer for size distribution detection.

### RNA-seq data processing and viral diversity measures

The quality of the Fastq files was assessed with FASTQC (Andrews, 2010) and MultiQC (Ewels et al., 2016) and pre-processed as paired reads with BBDuk (https://sourceforge.net/projects/bbmap/). Adapters were removed, the first ten 5’ nucleotides of each read were cut, and the 3’ end sequences with average quality below 10 were removed. Processed reads shorter than 60 nucleotides were discarded. The parameter values were set to: ktrim = r, k = 31, mink = 11, qtrim = r, trimq = 10, maq = 5, forcetrimleft = 10, and minlength = 60. The processed files were mapped to JUv1580_vlc or JUv2572_vlc as appropriated, with the BWA-MEM algorithm and default parameters (Li, 2013). Resulting SAM files were binarized and sorted with SAMtools (Danecek et al., 2021).

Viral variants were called with LoFreq (Wilm et al., 2012), only frequencies above 0.01 where considered. Nucleotide frequencies per position were obtained with pysamstats v1.1.2 (--type variation; https://hpc.nih.gov/apps/pysamstats.html). Normalized Shannon entropy per position was calculated as in Gregori et al. (2014) with the NormShannon method from QSutils v1.16.1 (Guerrero-Murillo and Gregori, 2023). Higher entropy values indicate more heterogeneous viral populations. Conversely, lower entropy indicates genetically homogeneous populations (Shannon, 1948; Gregory et al., 2016).

### Protein structure prediction

JUv1580_vlc RdRp structure was predicted using the standard parameters of ColabFold version 1.5.5 (Mirdita et al., 2022). For the structure of the CP, the updated PDB 4NWV (Guo et al., 2014) was used. For the structure of δ, PDB 5W82 (Guo et al., 2020) was used. Molecule visualizations were performed with ChimeraX (Meng et al., 2023).

### Statistical analysis

All statistical analyses mentioned in the previous sections were done with SPSS version 28.0.1.0 (IBM). In all cases, the sequential Bonferroni method was used for pairwise *post hoc* tests. Variance components associated to each factor in the GLMs described above were estimated by maximum likelihood. Graphs were generated using the “ggplot2” package version 3.4.0 (Wickham, 2009) in R version 4.1.1 within the RStudio development environment version 2021.09.0 Build 351.

## RESULTS

### Characterization of initial OrV inoculates

In order to perform experimental evolution with OrV strains JUv1580 and JUv2572, our first step was to characterize our particular isolates both genetically and phenotypically. Even though the original natural strains were previously characterized (Félix et al., 2011, Jiang et al., 2014, Frézal et al., 2019), since their isolation from nature they have infected an undetermined number of nematode cultures. During this culturing, new mutations could have fixed in both viral strains as a consequence of population bottlenecks or adaptation to laboratory growth conditions. Moreover, no sequence of JUv2572 is publicly available. To address this, we sequenced the genomes of our valencian OrV stocks, obtaining the genome of our JUv1580 (hereafter referred to as JUv1580_vlc; GenBank: XXXXXXX) and described for the first time a genome of JUv2572 (referred to as JUv2572_vlc; GenBank: XXXXXXX). In comparison with the reference genome of JUv1580 (Félix et al., 2011; GenBank: GCF_001402145.1), JUv1580_vlc has three synonymous mutations in RNA1 and four mutations in RNA2, one of them non-synonymous. When comparing the genome of JUv1580_vlc with JUv2572_vlc we observed 86 mutations in RNA1, 20 of which were non-synonymous, and 52 mutations in RNA2, ten of which were non-synonymous (Fig. 1A, Table S2).

**Figure 1.**
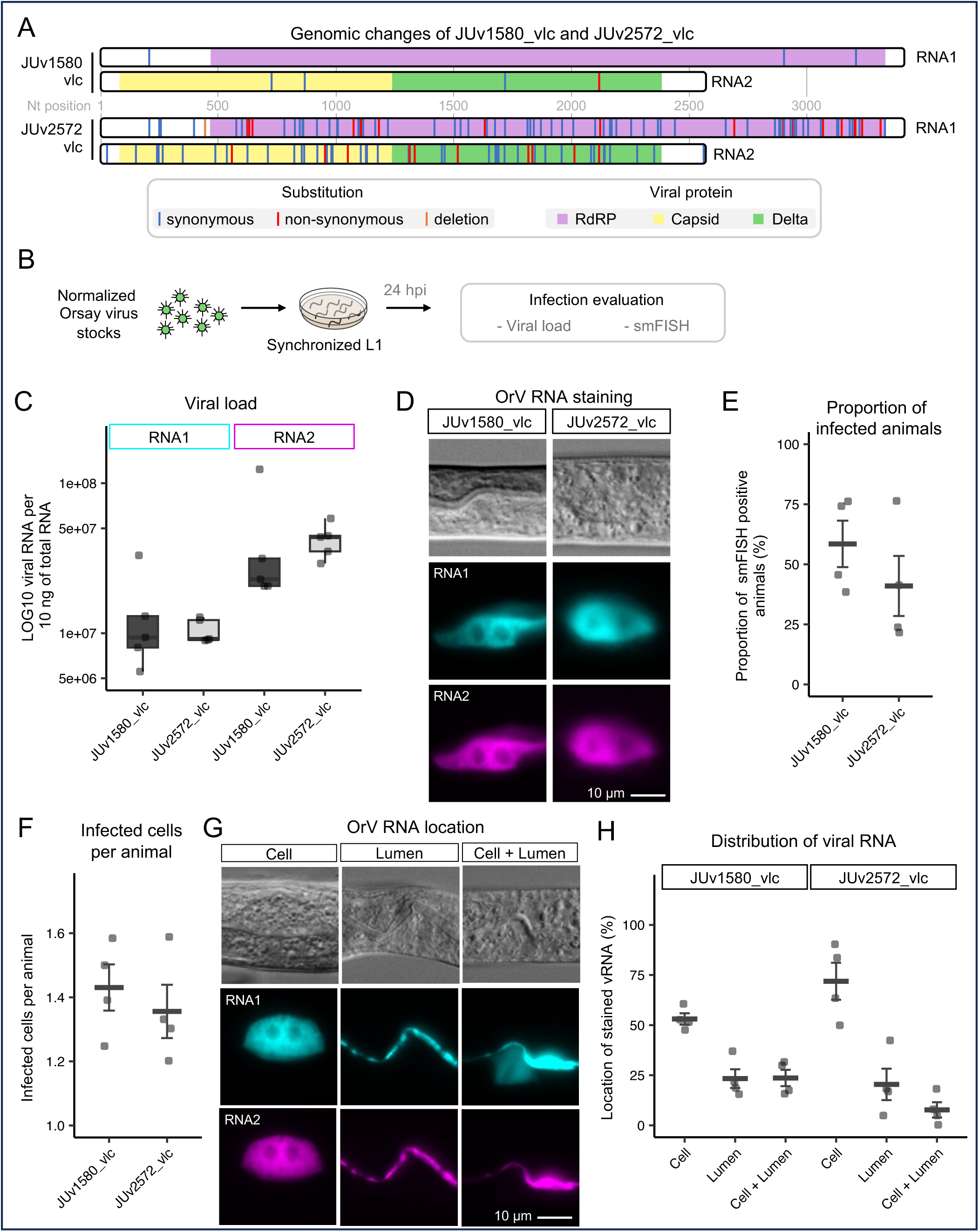
Comparison of the OrV strains used for experimental evolution. **(A)** Representation of the genomic differences of the strains JUv1580_vlc and JUv2572_vlc compared to the JUv1580 reference. **(B)** Experimental approach to evaluate the two strains. **(C)** Accumulation of RNA molecules (viral load) of the RNA1 (left panel, cyan) and RNA2 (right panel, magenta). **(D)** FISH staining of viral RNA for the JUv1580_vlc strain (left) and the JUv2572_vlc one (right). Top: DIC; middle: RNA1 in cyan; bottom: RNA2 in magenta. **(E)** Proportion of animals detected by FISH staining to be infected with JUv1580_vlc or JUv2572_vlc. **(F)** Average number of infected cells in four populations of 50 animals each. **(G)** Representative images of FISH-stained viral RNA present only in the cell (left), in the lumen (middle), or in both locations (right). Top: DIC; middle: RNA1 in cyan; bottom: RNA2 in magenta. **(H)** Proportion of the stained viral RNA located in intestinal cells, in the lumen, or in both locations. In (C) data are presented as median ±IQR, in the other panels as mean ±1 SE. Significance levels: *** *P* < 0.001; ** *P* < 0.01; * 0.01 ≤ *P* < 0.05; ^ns^ *P* ≥ 0.05.

To compare the infection phenotypes of our two viral strains, we prepared inocula of both strains with the same concentration of viral RNA. Synchronized L1 larvae of the ERT54 strain were inoculated and the infection was evaluated 24 hpi by RT-qPCR and by FISH staining of both viral RNAs. At 24 hpi, the number of copies (viral load) of RNA1 was the same for the two viral strains (GLM: *P* = 0.486) but both strains had more copies of RNA2 than of RNA1 (*P* < 0.001). The number of RNA2 molecules was 3.076 ±0.452 (mean ±1 SE) times higher than RNA1 molecules for JUv1580_vlc and 4.118 ±0.573 for JUv2572_vlc. We performed FISH staining of the viral RNA to examine the proportion of animals harboring infected cells, the number of infected cells and the location of the viral signal (Figs. 1D-G). We observed that the proportion of infected animals was 41.46% higher in animals inoculated with JUv1580_vlc in comparison with those inoculated with JUv2572_vlc (Fig. 1E, GLM: *P* < 0.001). The number of cells harboring viral RNAs in the infected animals was the same for both viral strains (Fig. 1F, *P* = 0.550). When examining the location of the viral RNAs, we observed viral RNAs within the intestinal cells, free in the lumen or in both locations (Fig. 1G). At 24 hpi, viral RNAs were predominantly found within intestinal cells upon infection with either viral strain (Fig. 1H, *P* < 0.001). However, we could detect a higher proportion of animals in the populations infected with JUv2572 showing viral RNA only in cells (Fig. 1H, *P* = 0.010). Altogether, these results indicate that RNA2 is present at higher levels than RNA1 and that JUv1580_vlc is able to infect more animals than JUv2572_vlc. However, JUv2572_vlc seems to have a higher replicative capacity at 24 hpi.

Finally, using inocula of both strains with the same concentration of viral RNA, we inoculated synchronized L1 ERT54 animals, and evaluated the infection 24 hpi or after three generations (Fig. 2A). We observed a 56.10% increase in animals activating the infection reporter *pals-5p::GFP* in populations inoculated with JUv1580_vlc (Fig. 2B, GLM: *P* < 0.001). Notably, after three generations the activation of *pals-5p::GFP* was higher than at 24 hpi (*P* < 0.001). The difference between the infection rate of animals that were directly inoculated and the infection rate of animals in which the virus was allowed to replicate for three generations was higher for JUv1580_vlc (57.14%) than for JUv2572_vlc (48.48%) (Fig. 2B, *P* = 0.026). These results highlight the importance of standardizing inoculation and analysis methods in order to compare viruses.

**Figure 2.**
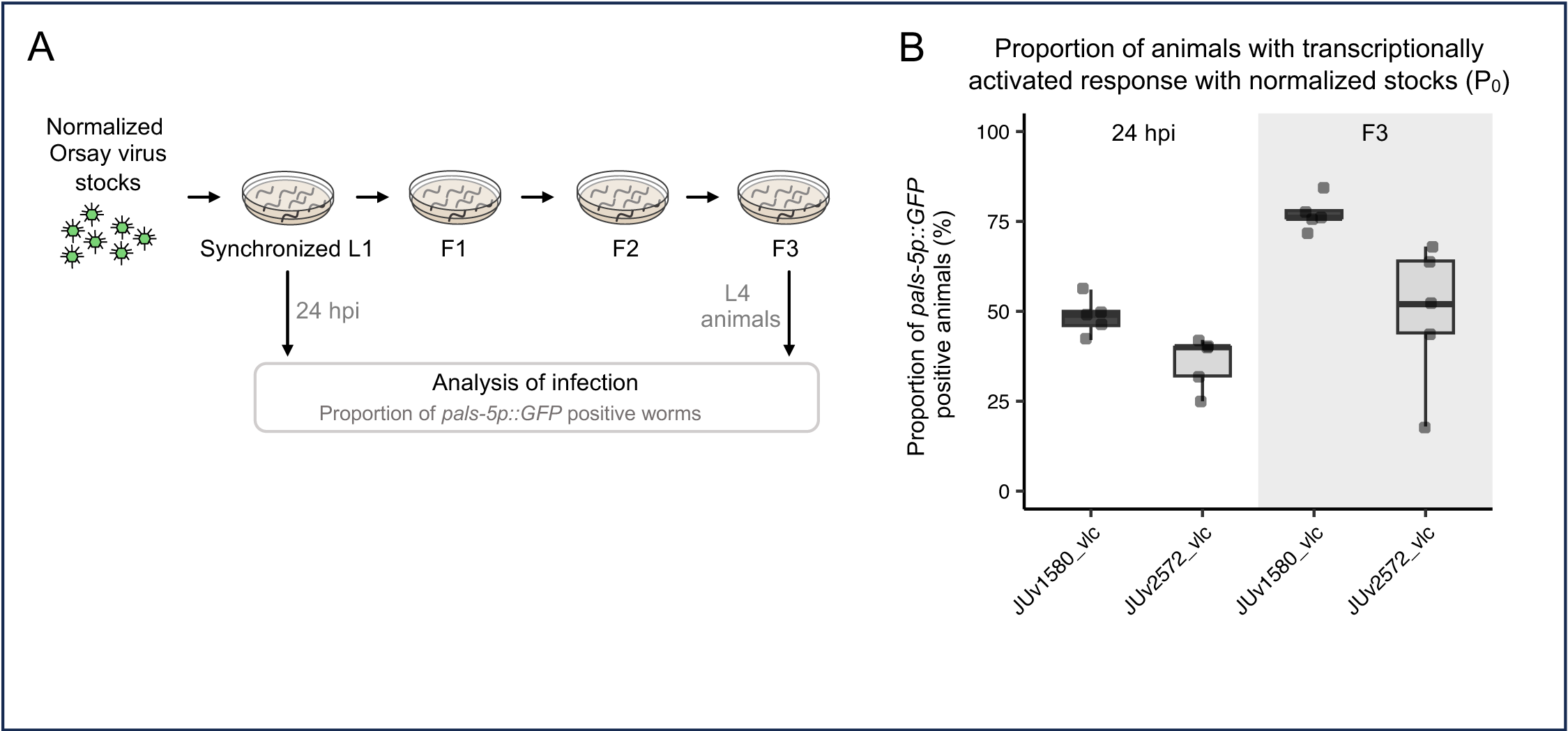
Characterization of ancestral OrV strains. **(A)** Experimental approach to evaluate the two strains. **(B)** Activation of the *pals-5p::GFP* reporter in ERT54 animals after being inoculated with normalized inocula of JUv1580_vlc or JUv2572_vlc. The infection was evaluated 24 hpi (left panel) or three generations after (right panel). Data are presented as median ±IQR.

### Changes in experimentally evolved OrV isolates

We performed a 10-passage experimental evolution experiment of the JUv1580_vlc and JUv2572_vlc viruses in ERT54 nematodes. The process was performed in triplicate for each virus, resulting in three experimentally evolved lineages per virus, referred to as R lineages. Moreover, another experimental block was performed for JUv2572_vlc in triplicate, obtaining another three experimentally evolved lineages of JUv2572, referred to as V lineages (Fig. 3A).

**Figure 3.**
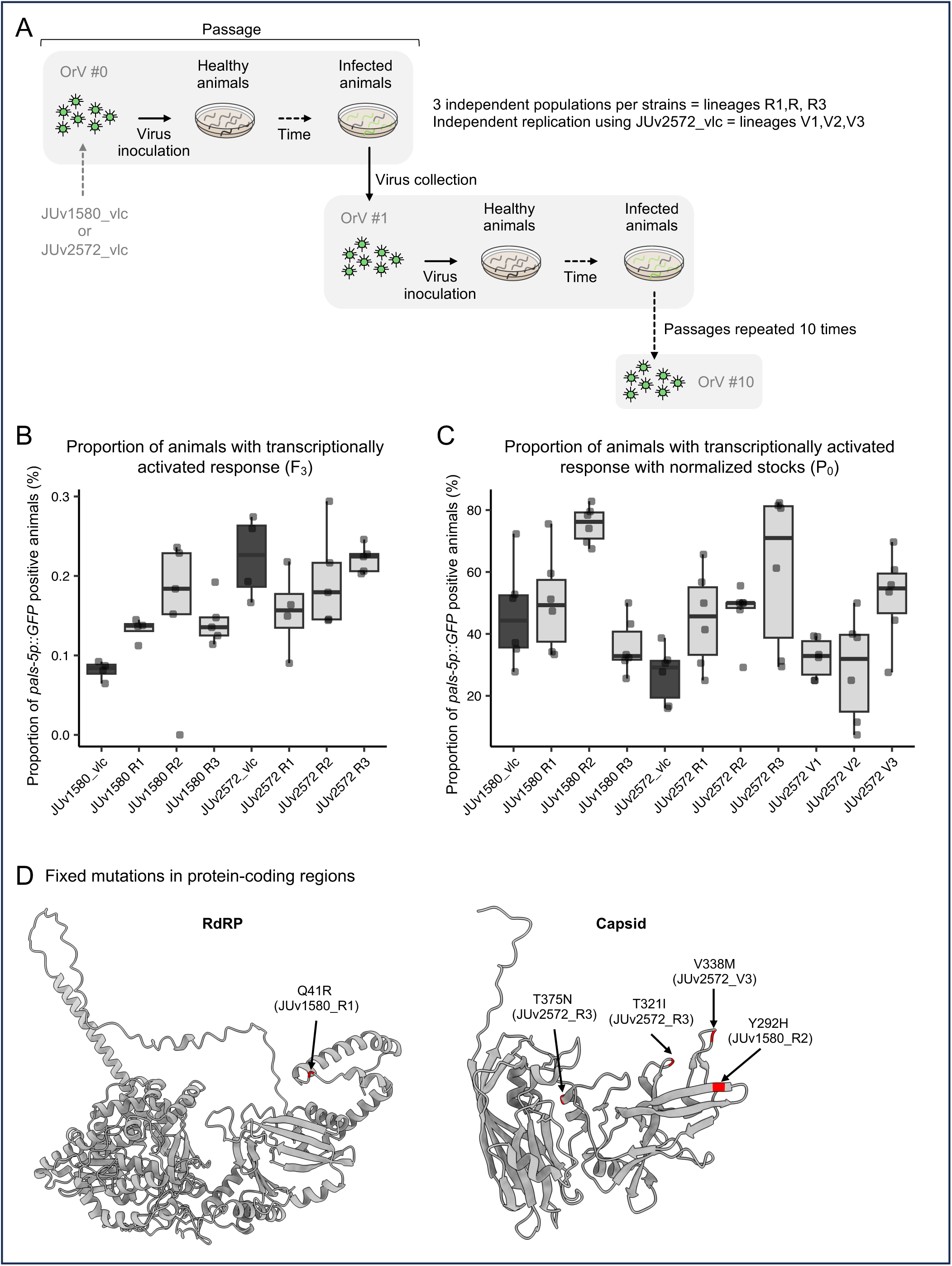
Characterization of experimentally evolved OrV strains. **(A)** Schematic representation of the experimental virus evolution (detailed in Methods). **(B)** Activation of the *pals-5p::GFP* reporter in ERT54 animals three generations (F_3_) after being inoculated with non-normalized inocula of ancestral and evolved strains. In this experiment, four replicate populations were evaluated per viral strain, with 100 animals assayed per replicate. **(C)** Activation of the *pals-5p::GFP* reporter in ERT54 animals 24 hpi with normalized inocula (P_0_) of ancestral and evolved strains. In this experiment, six replicate populations were evaluated per viral strain, with 33 ±6 animals (±1 SD) assayed per population. **(D)** Location of fixed amino acid changes in the evolved lineages. JUv1580_vlc RdRp structure was generated using AlphaFold and CP structure comes from PDB 4NWV (Guo et al., 2014). Data in all panels are presented as median ±IQR.

After the experimental evolution, we compared the infectivity of the evolved R and ancestral strains. Infectivity was measured by counting the proportion of adult animals activating the *pals5p::GFP* infection reporter at 24 hpi or after three generations post-inoculation. This approach was used to ensure that the final infection level was a function of the host-pathogen interaction over replication cycles rather than of the initial titer. We observed that, on average, linages evolved from the JUv2572_vlc showed no gain in infectivity when compared to their ancestral strain (Fig. 3B, *P* ≥ 0.448). However, the evolved linages derived from JUv1580_vlc had a greater infectivity than their ancestral strain (*P* ≤ 0.001) (Fig. 3B): JUv1580_R1 was 112.5% more infective, JUv1580_R2 was 125.0% more infective, and JUv1580_R3 showed a 75% increase. When compared with their ancestral, only the JUv2572_R1 lineage showed a significant infectivity change in comparison with its ancestral: a 30.4% reduction (*P* = 0.013).

We also evaluated the infectivity of the strains by normalizing the viral load of the inocula and quantifying the number of animals activating the *pals5p::GFP* reporter at 24 hpi. Using this approach, we observed a different pattern (Fig. 3C). The infectivity of evolved lineages derived from JUv1580_vlc was very variable: infectivity was higher than the ancestral for JUv1580_R2 (59.57% increase, *P* < 0.001), lower for JUv1580_R3 (23.40% decrease, *P* = 0.025), and equal for JUv1580_R1 (*P* = 0.422). For evolved lineages derived from JUv2572_vlc, all the R had higher infectivity than its ancestral lineage: JUv2572_R1 was 89.29% more infectious (*P* = 0.003). JUv2572_R2 was 60.71% more infectious (*P* = 0.001), and JUv2572_R3 had a 75% increase (*P* < 0.001). When assessing the infectivity of the V linages, only JUv2572_V3 had higher infectivity than the ancestral (89.29% increase, *P* < 0.001). Overall, V lineages showed 25.49% lower infectivity than R lineages (*P* < 0.001) (Fig. 3C).

### Test of genetic divergence versus evolutionary parallelism among linages

Did all nine independently derived genotypes exhibit the same suite of changes in their infectivity, or did they find alternative solutions to the challenge imposed by the host genotype and growth environment? At a first glance, Fig. 3C suggests substantial divergence in the derived genotypes, with statistically significant heterogeneity among lineages (Table 1). However, it is also relevant that most lineages showed an increase in infectivity when compared to the corresponding ancestor, with only JUv1580_R3 showing a reduction in infectivity. Following Vasi et al. (1994), the relative extent of genetic divergence versus phenotypic parallelism can be evaluated using the following index 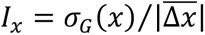, where *α_G_*(*x*) is the between-lineage genetic (additive) standard deviation for infectivity, and 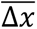 is the average change from the common ancestor. Notice that by using genetic standard deviation, both numerator and denominator have the same units. Hence, *I_x_* provides a measure of the average difference among independently derived lineages relative to the average evolutionary change from the ancestral state.

**Table 1.** Divergence *versus* parallelism in infectivity (*x*) for the nine independently derived lineages.

| OrV strain | Lineage | $x$ | $\overline{\Delta x}$ | $\sigma_G(x)$ | $I_x$ |
| --- | --- | --- | --- | --- | --- |
| JU1580_vlc | R1 | 0.511 ±0.033 | 0.068 ±0.116 | 5.954$\times 10^{-4}$ ±2.184$\times 10^{-7}$ | 2.303 ±3.932 |
| | R2 | 0.755 ±0.029 | | | |
| | R3 | 0.358 ±0.032 | | | |
| | Ancestral | 0.473 ±0.033 | | | |
| JUv2572_vlc | V1 | 0.320 ±0.035 | 0.168 ±0.048 | 6.439$\times 10^{-5}$ ±2.563$\times 10^{-9}$ | 0.535 ±0.155 |
| | V2 | 0.297 ±0.035 | | | |
| | V3 | 0.521 ±0.037 | | | |
| | R1 | 0.454 ±0.035 | | | |
| | R2 | 0.480 ±0.036 | | | |
| | R3 | 0.603 ±0.036 | | | |
| | Ancestral | 0.279 ±0.033 | | | |

In all cases, errors indicate $\pm 1$ SE.

Table 1 illustrates the effect of viral strain on the estimated *I_x_*. In the case of JUv1580_vlc-derived lineages, the index was greater than one, indicating that genetic differences among independently derived lineages are large relative to the average phenotypic change, *i.e*., the observed evolutionary difference in infectivity can mostly be explained by genetic differences among lineages. By contrast, in the case of JUv2572_vlc-derived lineages, the index was smaller than one, which indicates that the genetic differences among the independently derived viral lineages are small relative to the average change in infectivity from the ancestral state, *i.e*., non-heritable differences in infectivity had contributed to the observed phenotypic changes.

### OrV hotspots for nucleotide substitutions

Four of the evolved lineages (JUv1580_R1, JUv1580_R2, JUv2572_V3, and JUv2572_R3) were sequenced by high-throughput RNA-seq. Analysis of the evolved viral lineages revealed a low frequency of fixed mutations, with only three mutations fixed in each of the four lineages (Table S3). Out of these 12 different mutations, only five were non-synonymous substitutions: one in the RdRp of JUv1580_vlc_R1 (Q41R), one in the CP of JUv1580_vlc_R2 (Y292H), and three in the capsid of lineages derived from JUv2572_vlc: two in lineage R3 (T321I and T375N) and one in V3 (V338M) (Fig. 3D and Table S3). Of note, all of the mutations in CP are located within the P domain which forms the trimeric surface protrusions (Guo et al., 2014).

We next assessed the amount of genetic diversity present within the evolved populations by quantifying the per base normalized Shannon entropy. Higher entropy values indicate more heterogeneous viral populations, which contain intermediate-frequency variants (Shannon, 1948; Gregory et al., 2016). In contrast to the fixed differences, here we observed similar hotspots for nucleotide diversity in the four evolved lineages analyzed, independently of the ancestral strain they originated from (Fig. 4A). In RNA1, we observed hotspots between nucleotides 250-500 and 3100-3300. In RNA2, we found a hotspot in the first 20 nucleotides, and two others between nucleotides 400-500, within the S domain of the CP (Guo et al., 2014), and 1500-1600, from the end of the second alpha sheet to beta sheet 6 of the δ protein (Fig. 4A, B) (Guo et al., 2020).

**Figure 4.**
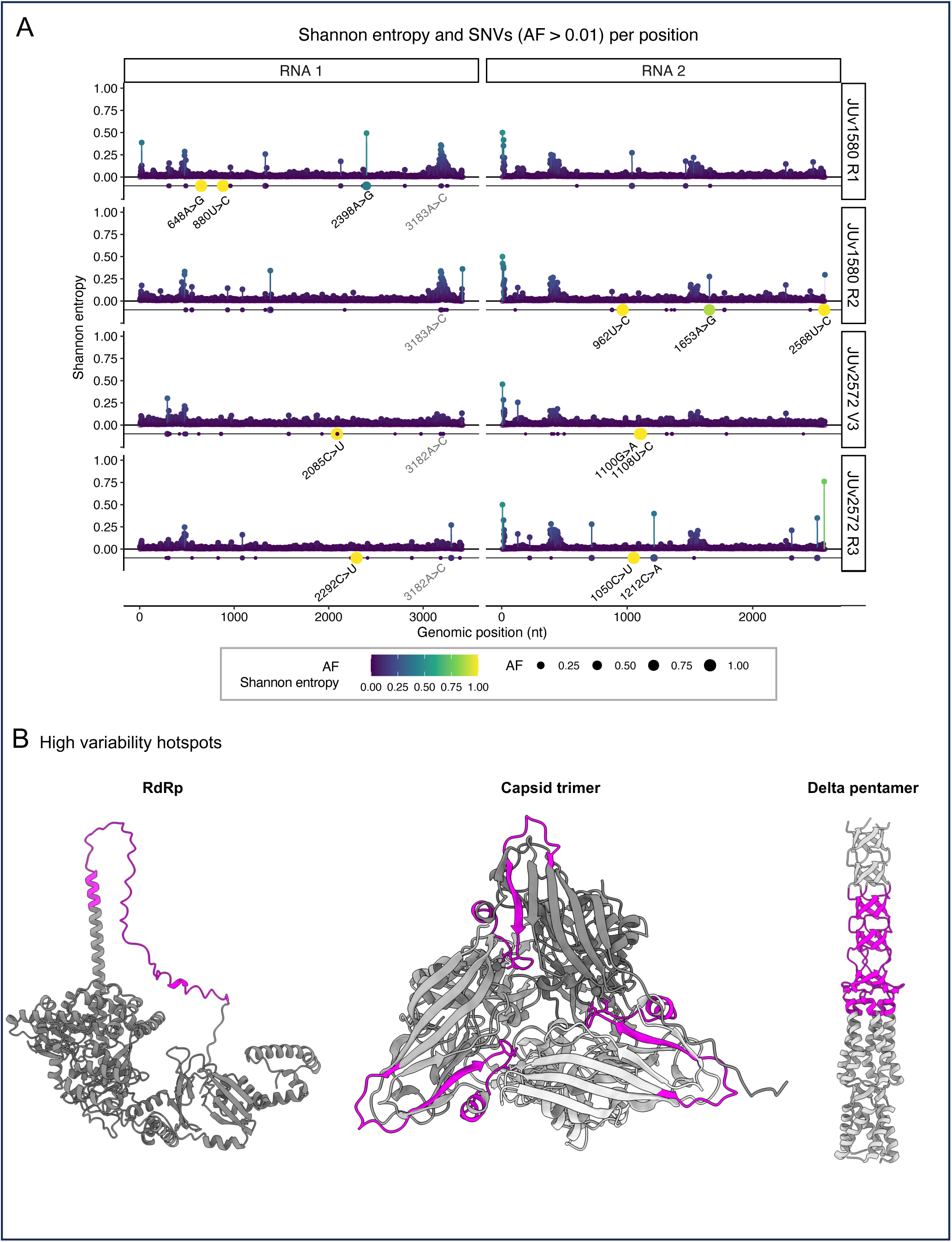
Population diversity of experimentally evolved strains. **(A)** Per-site normalized Shannon entropy for each RNA segment and viral strain SNVs with allele frequency (AF) > 0.01 are represented as dots in the line below each segment. Size and color display AF values. SNVs with and AF ≈ 1 are labeled in black. The convergent SNVs above the selected threshold, RNA 1: 3183A>C, is highlighted in grey. AF > 0.2 are labeled in black. The other common SNV (RNA 1: 302G>A) was not above the threshold in sample JUv1580_vlc RL2. **(B)** High variability hotspots (magenta) represented on the predicted structure of RdRp and PDB structures 4NWV and 5W82 of the CP trimer and δ pentamer (amino acids 1-110), respectively (Guo et al., 2014, Guo et al., 2020).

We could detect many other minor variants in the four genomes sequenced (Fig. 4A). These variants appeared at different frequencies, with the lower frequency detected confidently being 0.01 (Supplementary Table S4). Notably, two of the minor variants in RNA1 were detected in all samples: 302G>A and 3183A>C in JUv1580_vlc and its corresponding 3182A>C in JUv2572_vlc (Supplementary Table S5). These minor variants originated in parallel in the four lineages and are in the hotspot regions of the genomes.

## DISCUSSION

Experimental virus evolution can mimic natural processes (Elena and Sanjuán, 2007; Hajimorad et al., 2011). The availability of convenient pathosystems is key for experimental evolution. The discovery of OrV enables the study of viruses in *C. elegans*, an animal that has many advantages as model system: its small size enables work with large populations, its rapid life cycle enables us to perform experiments in a relatively short time, and its tolerance to freezing and thawing enables the preservation of samples (Corsi et al., 2015). In addition, this pathosystem is gaining diversity in the strains available, which enables a deeper understanding of the host-virus interactions (Shaw and Kennedy 2022). In *C. elegans* this applies to (*i*) the virus, as the efforts for isolating new strains and virus species are increasing (Frézal et al. 2029); (*ii*) the host, as there are large collections of natural isolates (Cook et al., 2017), genetically modified *Caenorhabditis* (Daul et al., 2019) and modified strains can be made using genome editing (Friedland et al., 2013); and (*iii*) the bacteria used to culture nematodes, as collections of the bacteria associated with *C. elegans* in nature are available (Berg et al., 2016; Dirksen et al., 2016; Samuel et al., 2016) and have been shown to modulate viral infections (Vassallo et al., 2023; González and Félix, 2024b). This rich diversity enables interesting future studies of co-evolution of the virus with its host and/or associated microbes.

We characterized here two strains of OrV. Both strains showed a genome formula with a clear dominance of RNA2 to RNA1 molecules. However, the proportion for each strain was slightly different, which could suggest a genome formula adaptation at the strain level. Indeed, viruses can adjust their genome formula for different reasons, such as to rapidly adapt to new environments (Zwart and Elena, 2020; Sicard et al., 2023; Bonnamy et al., 2024). One limitation here is that we only assessed these parameters at one time point, and the differences in RNAs proportion could also be due to different viral dynamics for both strains. Indeed, we observed different RNA localization patterns for both strains, which have been previously linked to different infection stages of OrV (Castiglioni et al., 2023). A more detailed analysis of the viral dynamics of these strains could shed light into the relevance of the genome formula.

Comparing viruses phenotypically is challenging. We propose here a normalization of the viral load present in the inoculum. A caveat of this approach is that it does not consider possible defective particles interfering with the infectivity of the inoculum. However, we do observe the same infectivity trends when inoculating with normalized stocks as when analyzing the infected animals after three generations (as proposed in Frézel et al., 2019): JUv1580_vlc appears to be more infectious than JUv2572_vlc. This results, which are in contrast to previous studies that also analyzed the infected worms three generations after inoculation (Frézal 2019), suggest that normalization of initial inoculum is relevant for subsequent analysis, even after three generations.

Another way of comparing viral strains is comparing their genomic sequences. The advent of high-throughput sequencing technologies, coupled with sophisticated computational analyses, has revolutionized our ability to genetic variability, highlighting the importance of minor variants within viral populations. This enhanced capability enables a more detailed characterization of the viral quasispecies, providing deeper insights into the composition and dynamics of viral populations. Such advancements are crucial for predicting the evolutionary trajectories of viruses and understanding their adaptability (Lauring and Andino, 2010). Altogether, refining the approach to compare strains is essential to describe differences in the infection phenotypes.

We have observed a certain degree of parallelism among the lineages evolved from the same ancestor. The parallelism was stronger for lineages derived from the JUv1580_vlc strain, with an excess of genetic variability for infectivity compared to the observed phenotypic change. The opposite situation was observed for lineages evolved from the JUv2572_vlc strain, with parallelism being mostly explained by non-genetic factors (*e.g*., contingency or chance). At odds with these observations, the amount of SNVs observed in the four sequenced lineages was not different among lineages derived from both strains. Furthermore, few fixed mutations were observed in our evolved OrV strains. The substitution rate –the rate at which mutations become fixed within a population– for a particular virus depends on several parameters, such as the environmental conditions and the mutation rate of that particular virus (Belshaw et al., 2011; Sanjuán, 2012). Given that RNA viruses are typically considered to have high mutation rates, the low number of fixed mutations we observed could be because our OrV strains have already adapted to replication in standard *C. elegans* laboratory conditions and had already exhausted most of beneficial substitutions. Alternatively, it is possible that our experimental conditions did not favor the accumulation of beneficial substitutions. The latter would usually result from (*i*) very low mutation rates, (*ii*) weak positive selection and strong genetic drift, which may lead to the loss of beneficial alleles before they reach a high frequency, and (*iii*) negative epistatic effects among beneficial mutations as would result from overlapping genes or multifunctional proteins that impose additional constraints to molecular evolution (Belshaw et al., 2008). Further analysis of OrV evolution and a more complete understanding of its genome would be necessary to clarify this apparent molecular evolution stasis.

## CONCLUSION

Experimental evolution of viruses can contribute to illuminate the complexities of evolutionary processes. *C. elegans* characteristics make it an interesting host for experimentally evolving viruses. Our work paves the ground for the use of the *C. elegans*-OrV pathosystem in future research on the evolution host-pathogens interactions.

## ACKNOWLEDGEMENTS

We thank Francisca de la Iglesia, Paula Agudo, and Aurélien Richaud for excellent technical support. We thank WormBase. Computations were performed on the HPC cluster Garnatxa at the Institute for Integrative Systems Biology (I2SysBio). This work was supported by grants PID2022-136912NB-I00 funded by MCIN/AEI/10.13039/501100011033 and by “ERDF a way of making Europe”, and CIPROM/2022/59 funded by Generalitat Valenciana to S.F.E. V.G.C. was supported by grant FJC2021-047264-I funded by MCIN/AEI/10.13039/501100011033 and by NextGenerationEU/PRTR. M.J.O.-U. was supported by grant FPU2019/05246 funded by MCIN/AEI/10.13039/501100011033 and by “ESF investing in your future”. R.G. was funded by EMBO (STF 8687 and Postdoctoral Fellowship ALTF 311-2021).

## REFERENCES

Andrews, S. (2010). FASTQC: A quality control tool for high through-put sequence data. http://www.bioinformatics.babraham.ac.uk/projects/fastqc/

Ashe, A., Bélicard, T., Le Pen, J., Sarkies, P., Frézal, L., Lehrbach, N.J., Félix, M.-A., Miska, E.A., 2013. A deletion polymorphism in the *Caenorhabditis elegans* RIG-I homolog disables viral RNA dicing and antiviral immunity. eLife 2, e00994.

Bakowski, M.A., Desjardins, C.A., Smelkinson, M.G., Dunbar, T.A., Lopez-Moyado, I.F., Rifkin, S.A., Cuomo, C.A., Troemel, E.R., 2014. Ubiquitin-mediated response to microsporidia and virus infection in *C. elegans*. PLoS Pathogens 10, e1004200. 10.1371/journal.ppat.1004200

Belshaw, R., Gardner, A., Rambaut, A., Pybus, O.G., 2008. Pacing a small cage: mutation and RNA viruses. Trends in Ecology and Evolution 23, 188–193. 10.1016/j.tree.2007.11.010

Belshaw, R., Sanjuán, R., Pybus, O.G., 2011. Viral mutation and substitution: units and levels. Current Opinion in Virology 1, 430–435. 10.1016/j.coviro.2011.08.004

Berg, M., Stenuit, B., Ho, J., Wang, A., Parke, C., Knight, M., Alvarez-Cohen, L., Shapira, M., 2016. Assembly of the *Caenorhabditis elegans* gut microbiota from diverse soil microbial environments. ISME Journal 10, 1998–2009. 10.1038/ismej.2015.253

Bonnamy, M., Brousse, A., Pirolles, E., Michalakis, Y., Blanc, S., 2024. The genome formula of a multipartite virus is regulated both at the individual segment and the segment group levels. PLoS Pathogens 20, e1011973. 10.1371/journal.ppat.1011973

Brenner, S., 1974. The genetics of *Caenorhabditis elegans*. Genetics 77, 71–94. 10.1093/genetics/77.1.71

Bull, J.J., Badgett, M.R., Wichman, H.A., Huelsenbeck, J.P., Hillis, D.M., Gulati, A., Ho, C., Molineux, I.J., 1997. Exceptional convergent evolution in a virus. Genetics 147, 1497–1507. 10.1093/genetics/147.4.1497

Castiglioni, V.G., Olmo-Uceda, M.J., Villena-Giménez, A., Muñoz-Sánchez, J.C., Elena, S.F., 2023. Story of an infection: viral dynamics and host responses in the *Caenorhabditis elegans*-Orsay virus pathosystem. bioRxiv. 10.1101/2023.10.31.564947

Cook, D.E., Zdraljevic, S., Roberts, J.P., Andersen, E.C., 2017. CeNDR, the *Caenorhabditis elegans* natural diversity resource. Nucleic Acids Research 45, D650–D657. 10.1093/nar/gkw893

Cornwall, D.H., Kubinak, J.L., Zachary, E., Stark, D.L., Seipel, D., Potts, W.K., 2018. Experimental manipulation of population-level MHC diversity controls pathogen virulence evolution in *Mus musculus*. Journal of Evolutionary Biology 31, 314–322. 10.1111/jeb.13225

Corsi, A.K., Wightman, B., Chalfie, M., 2015. A transparent window into biology: a primer on *Caenorhabditis elegans*. Genetics 200, 387–407. 10.1534/genetics.115.176099

Danecek, P., Bonfield, J.K., Liddle, J., Marshall, J., Ohan, V., Pollard, M., Whitwham, A., Keane, T., McCarthy, S.A., Davies, R.M., Li, H., 2021. Twelve years of SAMtools and BCFtools. GigaScience 10, giab008. 10.1093/gigascience/giab008

Daul, A.L., Andersen, E.C., Rougvie, A.E. 2019. The *Caenorhabditis* genetics center (CGC) and the C*aenorhabditis elegans* natural diversity resource. In The Biological Resources of Model Organisms (pp. 69–94). CRC Press.

Dirksen, P., Marsh, S.A., Braker, I., Heitland, N., Wagner, S., Nakad, R., Mader, S., Petersen, C., Kowallik, V., Rosenstiel, P., Félix, M.-A., Schulenburg, H., 2016. The native microbiome of the nematode *Caenorhabditis elegans*: gateway to a new host-microbiome model. BMC Biology 14, 38. 10.1186/s12915-016-0258-1

Domingo, E., Perales, C., 2019. Viral quasispecies. PLoS Genetics 15, e1008271. 10.1371/journal.pgen.1008271

Ebert, D., 1998. Experimental evolution of parasites. Science 282, 1432–1436. 10.1126/science.282.5393.1432

Elena, S.F., 2012. RNA virus genetic robustness: possible causes and some consequences. Current Opinion in Virology 2, 525–530. 10.1016/j.coviro.2012.06.008

Elena, S.F., 2017. Local adaptation of plant viruses: lessons from experimental evolution. Molecular Ecology 26, 1711–1719. 10.1111/mec.13836

Elena, S.F., González-Candelas, F., Novella, I.S., Duarte, E.A., Clarke, D.K., Domingo, E., Holland, J.J., Moya, A., 1996. Evolution of fitness in experimental populations of vesicular stomatitis virus. Genetics 142, 673–679. 10.1093/genetics/142.3.673

Elena, S.F., Lenski, R.E., 2003. Evolution experiments with microorganisms: the dynamics and genetic bases of adaptation. Nature Reviews Genetics 4, 457–469. 10.1038/nrg1088

Elena, S.F., Sanjuán, R., 2007. Virus evolution: insights from an experimental approach. Annual Reviews of Ecology Evolution and Systematics 38, 27–52. 10.1146/annurev.ecolsys.38.091206.095637

Ewels, P., Magnusson, M., Lundin, S., Käller, M., 2016. MultiQC: Summarize analysis results for multiple tools and samples in a single report. Bioinformatics 32, 3047–3048. https://doi. org/10.1093/bioinformatics/btw354

Félix, M.-A., Ashe, A., Piffaretti, J., Wu, G., Nuez, I., Bélicard, T., Jiang, Y., Zhao, G., Franz, C.J., Goldstein, L.D., Sanroman, M., Miska, E.A., Wang, D., 2011. Natural and experimental infection of *Caenorhabditis* nematodes by novel viruses related to nodaviruses. PLoS Biology 9, e1000586. 10.1371/journal.pbio.1000586

Félix, M.-A., Wang, D., 2019. Natural viruses of *Caenorhabditis* nematodes. Annual Reviews of Genetics 53, 313–326. 10.1146/annurev-genet-112618-043756

Frézal, L., Jung, H., Tahan, S., Wang, D., Félix, M.-A., 2019. Noda-like RNA viruses infecting *Caenorhabditis* nematodes: sympatry, diversity, and reassortment. Journal of Virology 93, e01170–19. 10.1128/JVI.01170-19

Friedland, A.E., Tzur, Y.B., Esvelt, K.M., Colaiácovo, M.P., Church, G.M., Calarco, J.A., 2013. Heritable genome editing in *C. elegans* via a CRISPR-Cas9 system. Nature Methods 10, 741–743. 10.1038/nmeth.2532

Garijo, R., Hernández-Alonso, P., Rivas, C., Diallo, J.-S., Sanjuán, R., 2014. Experimental evolution of an oncolytic vesicular stomatitis virus with increased selectivity for p53-deficient cells. PLoS ONE 9, e102365. 10.1371/journal.pone.0102365

González, R., Butković, A., Escaray, F.J., Martínez-Latorre, J., Melero, I., Pérez-Parets, E., Gómez-Cadenas, A., Carrasco, P., Elena, S.F., 2021. Plant virus evolution under strong drought conditions results in a transition from parasitism to mutualism. Proceedings of the National Academy of Sciences of the USA 118, e2020990118. 10.1073/pnas.2020990118

González, R., Félix, M.-A., 2024a. *Caenorhabditis elegans* immune responses to microsporidia and viruses. Developmental and Comparative Immunology 154, 105148. 10.1016/j.dci.2024.105148

González, R., Félix, M.-A., 2024b. Naturally-associated bacteria modulate Orsay virus infection of *Caenorhabditis elegans*. PLoS Pathogens 20, e1011947. 10.1371/journal.ppat.1011947

Grass, V., Hardy, E., Kobert, K., Talemi, S.R., Décembre, E., Guy, C., Markov, P.V., Kohl, A., Paris, M., Böckmann, A., Muñoz-González, S., Sherry, L., Höfer, T., Boussau, B., Dreux, M., 2022. Adaptation to host cell environment during experimental evolution of Zika virus. Communications Biology 5, 1115. 10.1038/s42003-022-03902-y

Gregori, J., Salicrú, M., Domingo, E., Sánchez, A., Esteban, J.I., Rodríguez-Frías, F., Quer, J., 2014. Inference with viral quasispecies diversity indices: clonal and NGS approaches. Bioinformatics 30, 1104–1111. doi: 10.1093/bioinformatics/btt768

Grubaugh, N.D., Smith, D.R., Brackney, D.E., Bosco-Lauth, A.M., Fauver, J.R., Campbell, C.L., Felix, T.A., Romo, H., Duggal, N.K., Dietrich, E.A., Eike, T., Beane, J.E., Bowen, R.A., Black, W.C., Brault, A.C., Ebel, G.D., 2015. Experimental evolution of an RNA virus in wild birds: evidence for host-dependent impacts on population structure and competitive fitness. PLoS Pathogens 11, e1004874. 10.1371/journal.ppat.1004874

Guerrero-Murillo, M., Gregori i Font, J., 2023. QSutils: quasispecies diversity, R package version 1.20.0, https://bioconductor.org/packages/QSutils

Guo Y.R., Hryc C.F., Jakana J., Jiang H., Wang D., Chiu W., Zhong W., Tao YJ., 2014. Crystal structure of a nematode-infecting virus. Proceedings of the National Academy of Sciences of the USA 111, 12781–12786. 10.1073/pnas.1407122111

Guo, Y. R., Hryc C.F., Jakana J., Tao YJ., 2020. Orsay Virus CP-δ Adopts a Novel β-Bracelet Structural Fold and Incorporates into Virions as a Head Fiber. Journal of Virology 94, e01560–20. 10.1073/pnas.1407122111

Hajimorad, M.R., Wen, R.-H., Eggenberger, A.L., Hill, J.H., Maroof, M.A.S., 2011. Experimental adaptation of an RNA virus mimics natural evolution. Journal of Virology 85, 2557–2564. 10.1128/JVI.01935-10

Jiang, H., Franz, C.J., Wu, G., Renshaw, H., Zhao, G., Firth, A.E., Wang, D., 2014. Orsay virus utilizes ribosomal frameshifting to express a novel protein that is incorporated into virions. Virology 450–451, 213-221. 10.1016/j.virol.2013.12.016

Jumper, J., Evans, R., Pritzel, A., et al. (2021). Highly accurate protein structure prediction with AlphaFold. Nature 596, 583–589. 10.1038/s41586-021-03819-2

Keleta, L., Ibricevic, A., Bovin, N.V., Brody, S.L., Brown, E.G., 2008. Experimental evolution of human influenza virus h3 hemagglutinin in the mouse lung identifies adaptive regions in HA1 and HA2. Journal of Virology 82, 11599–11608. 10.1128/JVI.01393-08

Kubinak, J.L., Ruff, J.S., Cornwall, D.H., Middlebrook, E.A., Hasenkrug, K.J., Potts, W.K., 2013. Experimental viral evolution reveals major histocompatibility complex polymorphisms as the primary host factors controlling pathogen adaptation and virulence. Genes and Immunity 14, 365–372. 10.1038/gene.2013.27

Lauring, A.S., Andino, R., 2010. Quasispecies theory and the behavior of RNA viruses. PLoS Pathogens 6, e1001005. 10.1371/journal.ppat.1001005

Lewis, J.A., Morran, L.T., 2022. Advantages of laboratory natural selection in the applied sciences. Journal of Evolutionary Biology 35, 5–22. 10.1111/jeb.13964

Lezcano, O.M., Fuhrmann, L., Ramakrishnan, G., Beerenwinkel, N., Huynen, M.A., Van Rij, R.P., 2023. Parallel evolution and enhanced virulence upon *in vivo* passage of an RNA virus in *Drosophila melanogaster*. Virus Evolution 9, vead074. 10.1093/ve/vead074

Li, H., 2013. Aligning sequence reads, clone sequences and assembly contigs with BWA-MEM. arXiv, http://arxiv.org/abs/1303.3997

Martineau, C.N., Kirienko, N.V., Pujol, N., 2021. Innate immunity in *C. elegans*. Current Topics in Developmental Biology 144, 309–351. 10.1016/bs.ctdb.2020.12.007

Martínez, J., Bruner-Montero, G., Arunkumar, R., Smith, S.C.L., Day, J.P., Longdon, B., Jiggins, F.M., 2019. Virus evolution in *Wolbachia-* infected *Drosophila*. Proceedings of the Royal Society B. 286, 20192117. 10.1098/rspb.2019.2117

McDonald, M.J., 2019. Microbial experimental evolution – a proving ground for evolutionary theory and a tool for discovery. EMBO Reports 20, e46992. 10.15252/embr.201846992

Meng, E.C., Goddard, T.D., Pettersen, E.F., Couch, G.S., Pearson, Z.J., Morris, J.H., Ferrin, T.E., 2023. UCSF ChimeraX : Tools for structure building and analysis. Protein Science 32, e4792. 10.1002/pro.4792

Meyer, J.R., Dobias, D.T., Weitz, J.S., Barrick, J.E., Quick, R.T., Lenski, R.E., 2012. Repeatability and contingency in the evolution of a key innovation in phage lambda. Science 335, 428–432. 10.1126/science.1214449

Mirdita, M., Schütze, K., Moriwaki, Y., Heo, L., Ovchinnikov, S., Steinegger, M., 2022. ColabFold: making protein folding accessible to all. Nature Methods 19, 679–682. 10.1038/s41592-022-01488-1

Mongelli, V., Lequime, S., Kousathanas, A., Gausson, V., Blanc, H., Nigg, J., Quintana-Murci, L., Elena, S.F., Saleh, M.-C., 2022. Innate immune pathways act synergistically to constrain RNA virus evolution in *Drosophila melanogaster*. Nature Ecology and Evolution 6, 565–578. 10.1038/s41559-022-01697-z

Nance, J., Frøkjær-Jensen, C., 2019. The *Caenorhabditis elegans* transgenic toolbox. Genetics 212, 959–990. 10.1534/genetics.119.301506

Navarro, R., Ambrós, S., Butković, A., Carrasco, J.L., González, R., Martínez, F., Wu, B., Elena, S.F., 2022. Defects in plant immunity modulate the rates and patterns of RNA virus evolution. Virus Evolution 8, veac059. 10.1093/ve/veac059

Pagán, I., Fraile, A., Fernandez-Fueyo, E., Montes, N., Alonso-Blanco, C., García-Arenal, F., 2010. *Arabidopsis thaliana* as a model for the study of plant–virus co-evolution. Philosophical Transactions of the Royal Society B 365, 1983–1995. 10.1098/rstb.2010.0062

Parker, D.M., Winkenbach, L.P., Parker, A., Boyson, S., Nishimura, E.O., 2021. Improved methods for single-molecule fluorescence *in situ* hybridization and immunofluorescence in *Caenorhabditis elegans* embryos. Current Protocols 1, e299. 10.1002/cpz1.299

Poole, D.S., Yú, S., Caì, Y., Dinis, J.M., Müller, M.A., Jordan, I., Friedrich, T.C., Kuhn, J.H., Mehle, A., 2014. Influenza A virus polymerase is a site for adaptive changes during experimental evolution in bat cells. Journal of Virology 88, 12572–12585. 10.1128/JVI.01857-14

Richaud, A., Frézal, L., Tahan, S., Jiang, H., Blatter, J.A., Zhao, G., Kaur, T., Wang, D., Félix, M.-A., 2019. Vertical transmission in *Caenorhabditis* nematodes of RNA molecules encoding a viral RNA-dependent RNA polymerase. Proceedings of the National Academy of Sciences of the USA 116, 24738–24747. 10.1073/pnas.1903903116

Samuel, B.S., Rowedder, H., Braendle, C., Félix, M.-A., Ruvkun, G., 2016. *Caenorhabditis elegans* responses to bacteria from its natural habitats. Proceedings of the National Academy of Sciences of the USA 113, E3941–E3949. 10.1073/pnas.1607183113

Sanjuán, R., 2012. From molecular genetics to phylodynamics: evolutionary relevance of mutation rates across viruses. PLoS Pathogens 8, e1002685. 10.1371/journal.ppat.1002685

Sanjuán, R., 2010. Mutational fitness effects in RNA and single-stranded DNA viruses: common patterns revealed by site-directed mutagenesis studies. Philosophical Transactions of the Royal Society B 365, 1975–1982. 10.1098/rstb.2010.0063

Schindelin, J., Arganda-Carreras, I., Frise, E., Kaynig, V., Longair, M., Pietzsch, T., Preibisch, S., Rueden, C., Saalfeld, S., Schmid, B., Tinevez, J.-Y., White, D.J., Hartenstein, V., Eliceiri, K., Tomancak, P., Cardona, A., 2012. FIJI: an open-source platform for biological-image analysis. Nature Methods 9, 676–682. 10.1038/nmeth.2019

Shannon, C.E., 1948. A Mathematical theory of communication. Bell System Technical Journal 27, 379–423. 10.1002/j.1538-7305.1948.tb01338.x

Shaw, C.L., Kennedy, D.A., 2022. Developing an empirical model for spillover and emergence: Orsay virus host range in *Caenorhabditis elegans*. Proceedings of the Royal Society B 289, 20221165. 10.1098/rspb.2022.1165

Sicard, A., Yvon, M., Timchenko, T., Gronenborn, B., Michalakis, Y., Gutierrez, S., Blanc, S., 2013. Gene copy number is differentially regulated in a multipartite virus. Nature Communications 4, 2248. 10.1038/ncomms3248

Solé, R.V., Elena, S.F., 2019. Viruses as complex adaptive systems. Princeton University Press, Princeton. 10.2307/j.ctv69tgmm

Stiernagle, T., 2006. Maintenance of *C. elegans*. WormBook. 10.1895/wormbook.1.101.1

Tran, T.D., Luallen, R.J., 2024. An organismal understanding of *C. elegans* innate immune responses, from pathogen recognition to multigenerational resistance. Seminars in Cell and Developmental Biology 154, 77–84. 10.1016/j.semcdb.2023.03.005

Tsanov, N., Samacoits, A., Chouaib, R., Traboulsi, A.-M., Gostan, T., Weber, C., Zimmer, C., Zibara, K., Walter, T., Peter, M., Bertrand, E., Mueller, F., 2016. smiFISH and FISH-quant – a flexible single RNA detection approach with super-resolution capability. Nucleic Acids Research 44, e165–e165. 10.1093/nar/gkw784

Vasi, F., Travisano, M., Lenski, R.E., 1994. Long-term experimental evolution in *Escherichia coli*. II. Changes in life-history traits during adaptation to a seasonal environment. American Naturalist 144, 432–456. 10.1086/285685

Vassallo, B.G., Scheidel, N., Fischer, S.E.J., Kim, D.H., 2023. Bacteria are a major determinant of Orsay virus transmission and infection in *Caenorhabditis elegans*. bioRxiv. 10.1101/2023.09.05.556377

Vignuzzi, M., López, C.B., 2019. Defective viral genomes are key drivers of the virus– host interaction. Nature Microbiology 4, 1075–1087. 10.1038/s41564-019-0465-y

Wichman, H.A., Badgett, M.R., Scott, L.A., Boulianne, C.M., Bull, J.J., 1999. Different trajectories of parallel evolution during viral adaptation. Science 285, 422–424. 10.1126/science.285.5426.422

Wickham, H. 2009. ggplot2: Elegant graphics for data analysis. (Springer, New York, NY). 10.1007/978-0-387-98141-3

Wilm, A., Aw, P.P.K., Bertrand, D., Yeo, G.H.T., Ong, S.H., Wong, C.H., Khor, C.C., Petric, R., Hibberd, M.L., Nagarajan, N., 2012. LoFreq: A sequence-quality aware, ultra-sensitive variant caller for uncovering cell-population heterogeneity from high-throughput sequencing datasets. Nucleic Acids Research 40, 11189–11201. 10.1093/nar/gks918

Yuan, W., Zhou, Y., Fan, Y., Tao, Y.J., Zhong, W., 2018. Orsay δ protein is required for nonlytic viral egress. Journal of Virology 92, e00745–18. 10.1128/JVI.00745-18

Zwart, M.P., Elena, S.F., 2020. Modeling multipartite virus evolution: the genome formula facilitates rapid adaptation to heterogeneous environments. Virus Evolution 6, veaa022. 10.1093/ve/veaa022

